# Emerging neurodevelopmental mechanisms in patient induced pluripotent stem cells-derived spheroids modelling *SCN1A* Dravet Syndrome

**DOI:** 10.1101/2024.05.09.593441

**Authors:** Cristiana Mattei, Miaomiao Mao, Sean Byars, Erlina Mohamed Syazwan, Megan Oliva, Timothy Karle, Kay Richards, Ingrid Scheffer, Steven Petrou, Snezana Maljevic

## Abstract

*SCN1A* encodes Naᵥ1.1, a voltage-gated sodium channel preferentially expressed in GABAergic interneurons, and it is the major cause of Dravet Syndrome (DS), a rare condition of developmental and epileptic encephalopathy (DEE). Among over 1000 DS mutations reported to date, almost all cause *SCN1A* loss-of function (LoF). A reduction in NaV1.1 function in inhibitory neurons would subsequently cause an over-excitation of glutamatergic neurons resulting in seizures, which are exacerbated by the use of sodium channel blocking common anti-seizure medications (ASM). In this study we generated and assessed 3D spheroids enriched with GABAergic neurons from *SCN1A* DS patient to establish a 3D human-derived DS model. To investigate developmental disruptions in DS pathophysiology we profiled the transcriptome of patient-derived spheroids and subsequently, tested the capability of this 3D *in vitro* model to reveal the cellular mechanisms of DS and predict drug response. In summary, our patient iPSC-derived neuronal model of *SCN1A* DS revealed a profound dysregulation of developmental processes which correlated with functional disruption in GABAergic neurons and predicted response to fenfluramine, an ASM increasingly used for the treatment of DS.

## Introduction

Dravet syndrome (DS) is a severe developmental and epileptic encephalopathy with onset in infancy. It is a rare condition with a global prevalence ranging from one in 15,500 to one in 40,000 live births (1–3). In the first year of life, children present with drug-resistant epilepsy with frequent disabling seizures despite polypharmacy. Development prior to the seizure onset seems normal, but a slowing or regression is seen around the age of 2 years and all children develop, to variable degrees, an intellectual disability (4). DS associates with comorbidities, like behavioural problems, motor and sleep disruptions, having a major impact on the quality of life of the patients and their families (5). Additionally, patients with DS have an elevated mortality risk due to status epilepticus and SUDEP (Sudden Unexpected Death in People with Epilepsy) (6). DS is primarily caused by *de novo* loss-of function (LoF) variants in *SCN1A*. Mutations in other genes have been described in DS patients but are rare in patients with a core phenotype (7). *SCN1A* encodes the voltage-gated sodium channel Nav1.1, which is preferentially expressed in GABAergic interneurons and heterozygous LoF mutations lead to an haploinsufficiency. *SCN1A* pathogenic variants have been hypothesized to result in the disinhibition of excitatory neurons and a subsequent increase in the susceptibility to seizures and developmental impairments (8). However, gain-of-function (GoF) pathogenic variants in *SCN1A* have also been found in patients with early onset developmental and epileptic encephalopathies (DEEs) displaying clinical features distinct from DS. The underlying pathomechanism caused by *SCN1A* GoF variants is less clear, though computational modelling has suggested that an increase in interneuron activity could lead to network instability via changes in synaptic plasticity (9).

To date, there are no effective treatments available to ameliorate both epilepsy and developmental deficits in DS patients. While traditional anti-seizure medications (ASM) in the sodium channel blocker category (e.g. carbamazepine and phenytoin) are effective for treating GoF *SCN1A*/DEE patients, they tend to worsen seizures in LoF *SCN1A*/DS patients (10). However, a partial response can sometimes be achieved in DS patients using ASM with known mechanisms of action modulating neurotransmission (e.g. fenfluramine and stiripentol) (11).

Despite several DS mouse models being developed along with an exiguous number of *in vitro* studies (8)(12–18), the DS pathophysiology is not fully understood. Therefore, advancing our knowledge of the DS pathomechanisms in the human biology context would be beneficial to progress towards the development of effective therapeutic approaches. To this end, human induced pluripotent stem cell (iPSCs) technology with more recently developed 3D models, represents an unprecedented advance in studying human neurodevelopmental disorders.

In this study, we aimed to establish a 3D *in vitro* model of DS by the utilization of DS patient-derived iPS cell lines and isogenic control. Our approach is based on the region-specific spheroid technology (19,20) for the development of a robust differentiation protocol together with methodologies for unravelling DS pathophysiology and predicting efficacy of ASM.

## Materials and methods

### Generation and maintenance of iPSC lines and isogenic control from T1722N patient fibroblasts

Two distinct iPSC clones, 11 and 13, from patient fibroblasts and one clone of corresponding corrected cell line were generated by the Stem Cell Gene Editing Facility (MCRI, Victoria, Australia). Clonal iPSC populations were derived following introduction of episomal reprogramming vectors into patient fibroblasts using the Neon transfection system (1400 V, 20 ms, 2 pulses). Gene-editing factors were introduced into a single iPSC clone (clone 11) using the Neon transfection system (1100 V, 30 ms, 1 pulse). Transfected cells were plated over 2 wells of a Matrigel (cat #356234, Corning) coated 6-well dish in E8 medium (cat #A1517001, ThermoFisher) supplemented with 10 uM Y-27632 (Y27, cat #ab120129, Abcam). Medium changes were performed every day (without Y-27) and individual iPSC colonies were isolated and expanded. To identify targeted clones, colonies were screened by allele-specific PCR (using primers SCN1A_5165_corF: GTCTATTCCAAATTACAACCAGC and SCN1A_5165_scR: ATAGGAAGGTGGACAAGCTGC). Following multiple rounds of subcloning, a single corrected colony was selected and expanded. Subsequently, one corrected iPSC line and two patient iPSC lines were maintained onto 5µg/ml vitronectin (cat #A14700, ThermoFisher)-coated plates and passaged using 0.5 mM EDTA (cat #15575-038, Life Technologies). All cell lines progressed to quality control assessment.

### Quality control of iPSC lines

All cell lines used in this study underwent quality control tests prior to their use in experiments. Presence of mycoplasma in culture was assessed using the MicoAlert assay control test according to the manufacturer’s instructions (cat #LT07-518, Lonza). Sanger sequencing was performed to verify the presence of pathogenic variant in patient cell lines (clones 11 and 13) and its correction in isogenic control cell line, respectively. Genomic DNA was extracted from cell pellets using Dneasy Blood and Tissue kit (cat #69504, Qiagen) and amplicons of interest were amplified by PCR reaction (FW: ttcctagtcatgttcatctacgcc, RV: attagggtcacagtcgggtg). PCR products were cleaned up using QIAquick PCR Purification kit (cat #28106, Qiagen) and submitted to the Australian Genome Research Facility (AGRF) for sequencing. To assess karyotype of all the cell lines, cell pellets were submitted to the Victorian Clinical Genetics Services (VCGS) for Illumina Infinium GSA-24 v3.0 array. Stem cell pluripotency was assessed by immunostaining to verify the expression of pluripotency markers (see immunocytochemistry section), whereas to verify the differentiation potential, we used the STEMdiff trilineage differentiation assay according to the manufacturer’s instructions (cat #5230; Stem Cell Technologies) (see RT-qPCR section).

### Derivation of Subpullium spheroids

Subpullium Spheroids (SSs) were generated using a differentiation protocol adapted from (19)(20). Procedure for DIV 0 seeding and preparation of differentiation media, CDMI, CDMII, CDMIII and CDMIV were described by (20) (of note, Matrigel was not added to CDM and SSs were kept onto orbital shaker till termination of culture). To initiate the generation of SSs (DIV 0), iPSCs were lifted by using 0.5 mM EDTA and transferred into ultralow attachment 96-well plates (cat #7007, Corning) with CDMI medium supplemented with the ROCK inhibitor Y27, the SMAD inhibitor SB-431542 (SB, 10 µM, cat #72234, StemCell Technologies), the BMP inhibitor Dorsomorphin (DM, 3 µM, cat #P5499, Sigma Aldrich), and the WNT pathway inhibitor IWR-1 (3 µM, cat #681669, Millipore, Sigma-Aldrich). On DIV 2, 4, 6 and 9, medium was replaced with fresh supplemented CDMI (as per DIV 0) without Y27. On DIV 12, SSs were fed with CDMI supplemented with the SHH agonist, SAG (100 nM, cat #S7779, Selleckchem), IWR-1 (5 µM), recombinant human FGF (20 ng/mL, cat #233-FB, R&D), and recombinant human EGF (20 ng/mL, cat #236-EG, R&D). On DIV 15, fresh medium was given to SSs as per DIV 12 with addition of Allopregnanolone (AlloP, 100 nM, cat #16930, Cayman Chemical). On DIV 18, spheroids were transferred to untreated plates (cat #LBS60001X, Thermo Fisher Scientific) onto the orbital shaker (cat #NB-T101SRC, N-Biotek, Seoul, South Korea) and fed with CDMII medium supplemented as per DIV 15. Same procedure was repeated on DIV 21 whereas on DIV 24 and every third day till DIV 33, CDMII was supplemented with BDNF (20 ng/mL, cat #AF45002, Lonza), NT-3 (20 ng/mL, cat #45003, Peprotech), DHA (10 µM, cat #D2534, Sigma Aldrich), cAMP (100 µM, cat #D0627, Sigma Aldrich), and ascorbic acid (AA, 200 µM, cat #32144823, Novachem-Wako). On DIV 35 and 38, same supplements were maintained but CDMII was replaced by CDMIII. Subsequent medium change was performed on DIV 42 and 46 with unsupplemented CDMIII. On DIV 49 onwards, SSs were fed with unsupplemented CDMIV and medium change was performed every 3-4 days.

### RT-qPCR

Extraction of RNA samples and set up of RT-qPCR reactions were performed as previously described (21). Immunoreactivity. To obtain relative expression levels of each transcript we used single-plexing with the endogenous control HMBS. The specific probes (ThermoFisher) that have been used are as follow: BRACHYURY (Hs00610080_m1), CXCR4 (Hs00976734_m1), FOXG1 (Hs01850784_s1), HMBS (Hs00609297_m1), NKX2-1 (Hs00968940_m1), OTX2 (Hs00222238_m1), PAX6 (Hs01088114_m1), SOX17 (Hs00751752_s1), TBX6 (Hs00365539_m1), and TUBB3 (Hs00964962_g1). Data are shown as mean ± SD of n = 3 technical replicates, unless otherwise stated in figure legends. Statistical analysis was performed using GraphPad Prism 8 software. For the two independent sample comparison, the statistical significance was determined using the Holm–Sidak T-test and alpha=5.000%.

### Immunocytochemistry

Cells were fixed and stained as previously described (21). Immunoreactivity was tested against NANOG (#ab21624, Abcam, 1:100) and TRA-1-60(R) (#ab16288, Abcam, 1:100), The secondary antibodies were Alexa Fluor-conjugated 488 anti-Mouse (#A11001, Life Technologies) and Alexa Fluor-conjugated 594 anti-Rabbit (#A21207, Life Technologies). Nuclei were visualized using 4,6-diamidino-2-phenylindole, dihydrochloride (DAPI) counterstain (1 mg/mL; #D9452-50MG, Sigma-Aldrich). Images were acquired using a Zeiss 780 confocal microscope.

### Bulk RNA-sequencing and pathway analysis

To identify genes and pathways that may be dysregulated in DS SCN1A T1722N variant, mRNA sequencing of patient and control-derived SSs at DIV 35-40, 75-80 and 115-120 was performed. A total of n ≥ 2 biological replicates per timepoint was collected with multiple spheroids pooled together for each replicate. Samples were processed and analysed as described by (21).

### Whole-cell patch clamping

Patient and control-derived SSs at DIV144-168 were cut into 300 µm slices on a vibratome (VT1200S, Leica) in ice-cold cutting artificial cerebrospinal fluid (ACSF) containing (in mM) 125 mM Choline-Cl, 2.5 mM KCl, 0.4 mM CaCl2, 6 mM MgCl2, 1.25 mM NaH2PO4, 26 mM NaHCO3, and 20 mM D-glucose saturated with carbogen (95% oxygen and 5% carbon dioxide). Slices were incubated in recording ACSF consisting of 125 mM NaCl, 2.5 mM KCl, 2 mM CaCl2, 2 mM MgCl2, 1.25 mM NaH2PO4, 26 mM NaHCO3, and 10 mM D-glucose saturated with carbogen for 30 minutes at 32°C before returning to room temperature. They were recovered for at least one hour after sectioning before recordings were performed in a recording chamber at room temperature on an upright microscope (Slicescope Pro 1000, Scientifica). Slices were perfused with recording ACSF saturated with carbogen with a flow rate of 3 mL/min. Cells were identified with a 40x objective lens (Olympus) equipped with infrared-oblique illumination and patched with pipettes of 3 to 6 MΩ (#30-0060, Warner Instruments) made with a laser puller (P-2000, Sutter Instruments). The pipette solution contained (in mM): 130 K-gluconate, 10 D-glucose, 6 KCl, 5 EGTA, 5 HEPES, 4 NaCl, 2 MgATP, and 0.3 GTP-tris salt (pH 7.3 with KOH, 290 mOsm). Data were recorded with a Multiclamp 700B amplifier controlled by Clampex 10.6/DigiData 1440 acquisition systems (all from Molecular Devices) at a sampling rate of 10 kHz. Resting membrane potentials and spontaneous action potentials (AP) were recorded using I=0 gap-free protocol for > 30 seconds. Passive and active membrane properties were measured from voltage responses generated using a current clamp protocol in which the cell was held at -70 mV and injected with current from -10 to up to +35 pA for 1 sec. A calculated liquid junction potential of -12.2 mV was not corrected in the reported values. For voltage clamp recordings, cells were held at -70 mV and stepped between –70 mV and +40 mV for 1 sec before returning to –70 mV. Rs compensation was adjusted on MultiClamp at 1.02 kHz and 60%/80% for correction and prediction, respectively. For spontaneous postsynaptic current recordings, cells were held at –70 or +40 mV and recorded for five minutes. Cells with access resistance of >25 MΩ were excluded from analysis. Analyses were performed using Clampfit 10.6 (Molecular Devices, USA). Voltage clamp recordings were low pass filtered at 1 kHz offline for analysis. GraphPad Prism 9 (GraphPad, USA) was used for graphing and statistical analyses. Data contain recordings from at least three independent differentiations with the patient and isogenic control cell lines generated and recorded in parallel for each differentiation.

### Two-photon calcium imaging

Patient and isogenic control-derived SSs were loaded with 5 µM Calbryte^TM^ 520 AM (AAT Bioquest) for at least three hours protected from light prior to imaging. Briefly, thawed Calbryte^TM^ AM dye powder (50 µg) was dissolved in 20.25 µL of dimethylsulfoxide (DMSO) to make a 2 mM stock solution. A 10 µM 2X working solution of Calbryte^TM^ AM dye was prepared in sterile BrainPhys^TM^ Imaging Optimized Medium (BPI, STEMCELL Technologies) containing 0.08% pluronic F-127 (#20050, Sigma-Aldrich). The 2X working solution was diluted to make the final working solution in a 1.7 mL Eppendorf tube in which individual spheroids were incubated. Stock solutions of Calbryte^TM^ AM dye was kept in the fridge protected from light for up to 2 days.

Individual spheroids were mounted in a custom-made perplex perfusion dish and coverslipped. The dish was secured onto the imaging setup and the solution within was kept at 37 °C using a temperature controller (Warner Instrument). Two-photon (2P) Ca^2+^ imaging was performed with a 25x water immersion objective (Olympus XLPlan N 25X, NA = 1.05, correction collar = 0.17) on a 2P microscope built inhouse. A 100-femtosecond (fs) pulsed laser (Insight Spectra Physics) was tuned to 920nm to excite Calbryte^TM^ 520 AM. For each spheroid, data were collected from two distinct locations (Field of view = 275 µm x 275 µm) during baseline and 500 µM fenfluramine perfusion (±-fenfluramine hydrochloride, Novachem). Perfusion of fenfluramine was carried out at 1 mL/min and recordings were started 5 minutes after the beginning of perfusion. Each recording session contained 10000 frames sampled at 30 Hz. The base imaging solution used was BPI.

## Results

### SCN1A T1722N derived iPSCs differentiate into 3D neuronal cultures and show similar SCNxA gene expression

In this study we recruited a female individual diagnosed with DS and carrying a *de novo SCN1A* pathogenic variant T1722N (c.5165C>A) in heterozygosis. Position of the missense mutation T1722 is in the sixth membrane-spanning segment of Domain 4 of Nav1.1 (Figure S1A).

We derived two distinct iPSC lines, clones 11 and 13, from patient fibroblasts and one clone of corresponding corrected cell line. Cells in culture displayed typical morphological features of pluripotent cells with no sign of differentiation (Figure S1B, bottom). During expansion and entire duration of the study, cells were kept mycoplasma-free and karyotypically normal. The presence of the mutation in the patient derived iPSC lines and the correction in the isogenic control line was confirmed by Sanger sequencing (Figure S1B, top). The cell pluripotency stage was verified by immunofluorescence reactivity to pluripotent markers NANOG and TRA160R (Figure S1C, left). All cell lines displayed a normal karyotype (Figure S1C, right) and similar trilineage differentiation potential (Figure S1D).

To investigate how *SCN1A* T1722N genotype may impact human corticogenesis in DS, we have employed a 3D differentiation protocol giving arise to brain region-specific spheroids (19). Given the predominant expression of *SCN1A* in inhibitory cortical neurons (22), patient and control iPSCs were differentiated into subpullium-like spheroids (SSs) which were shown to resemble the ventral forebrain and are enriched with inhibitory neurons and their precursors (19). Our differentiation strategy consisted of three main phases, induction, expansion and maturation, driven by a strict modulation of specific signalling pathways (Figure 1A). Both patient and control iPSCs were capable of differentiating into SSs, showing a similar developmental trajectory over initial and later stages of differentiation (Figure 1B). The bulk RNA sequencing analysis revealed a substantial expression (>50 RPMK) of several ventral forebrain (or subpullium) markers at DIV 35-40 in both patient and control SSs. Conversely, dorsal forebrain markers displayed an overall scarce expression confirming the ventral specification of SSs (Figure 1B, top). At a later stage of differentiation, DIV 115-120, the GAD1+ population is the most represented with a predominant calbindin (CALB2) and somatostatin (SST) identity (Figure 1B, bottom).

**Figure 1.**
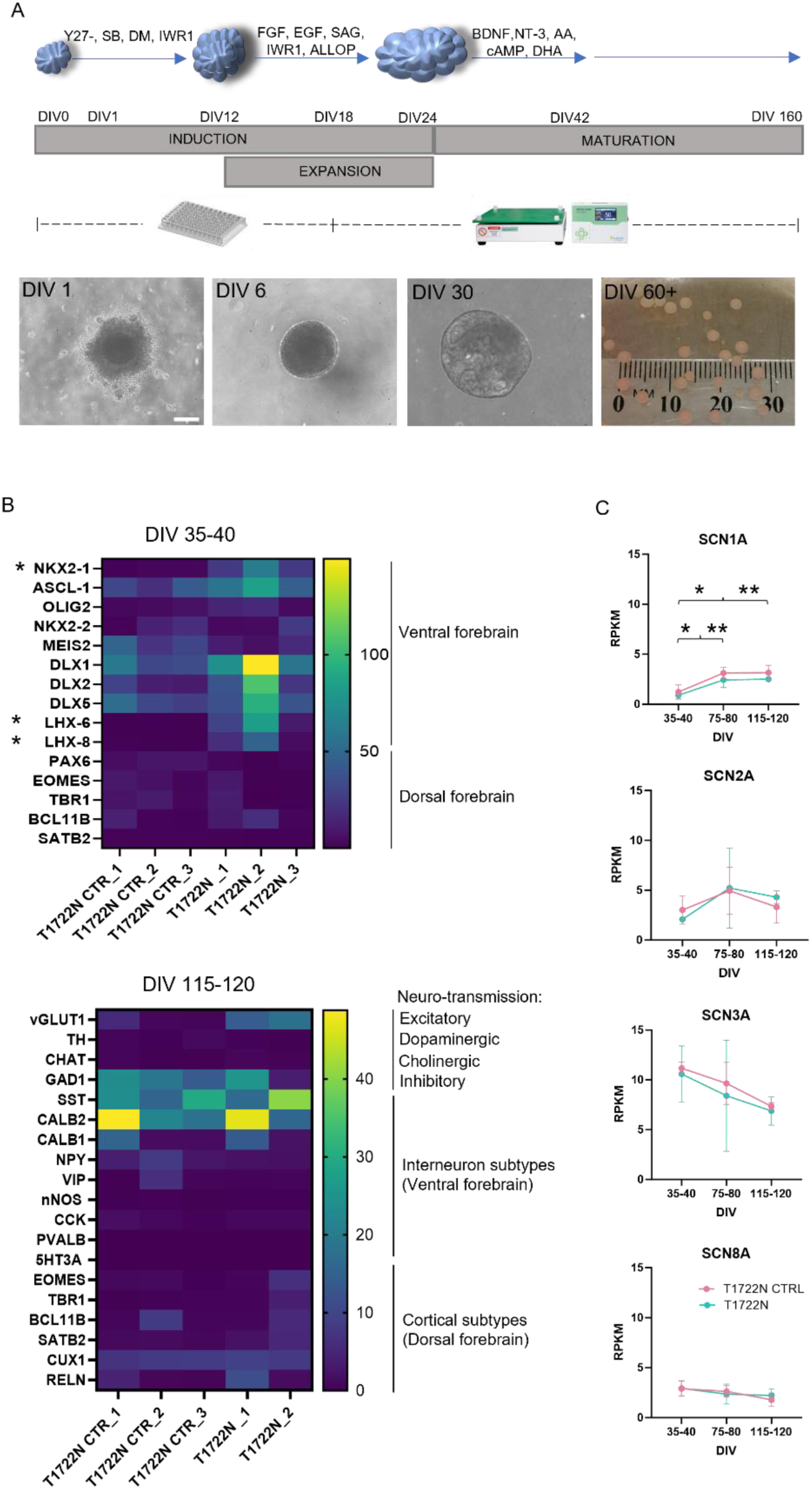
Derivation and characterization of SCN1A T1722N SSs. (A) Schematic of differentiation protocol (top) and representative bright field images of SSs at different timepoints of differentiation (bottom). (B) Expression of developmental markers at DIV 35-40 and 115-120 assessed by bulk RNA-sequencing (* FDR *q*<0.05). (C) Expression of *SCN1A*, *SCN2A*, *SCN3A*, and *SCN8A* at DIV 35-40, 75-80 and 115-120 in *SCN1A* T1722N patient and isogenic control-derived SSs assessed by bulk RNA-sequencing (CTRL DIV35-40 vs CTRL DIV75-80, **p=0.0077; CTRL DIV35-40 vs CTRL DIV115-120, **p=0.0069; T1722N DIV35-40 vs T1722N DIV75-80, *p=0.0436; T1722N DIV35-40 vs T1722N DIV115-120, *p=0.0353). Data presented in (C) are RPKM mean ± SD, from *n* = 3 biological replicates. Two-way ANOVA with multiple comparisons was used to assess statistical significance (*p*<0.05). Scale bars, 50 µm.

Expectedly, no differences in the expression of crucial differentiation markers at DIV 35-40 were found between the two patient cell lines, clones 11 and 13 (Figure S2A). Therefore, for the generation of subsequent datasets, only patient clone 13 was used and referred to as “T1722N”.

To assess the effect of T1722N mutation on *SCN1A* expression in this model, we next interrogate the *SCN1A* mRNA expression in SSs. This revealed an increased *SCN1A* expression over time with a statistically significant positive slope between DIV 35-40 and 75-80 in both genotypes. Expectedly, no differences were found between control and T1722N SSs, demonstrating that the missense mutation T1722N does not impact the levels of mRNA expression (Figure 1C). Other *SCN* genes found in the developing cortex, *SCN2A*, *SCN3A* and *SCN8A,* displayed similar expression profile in SSs between the two genotypes.

The capability of differentiating both patient and control iPSC lines into SSs posed the basis for our subsequent analysis aimed to interrogate pathomechanisms underlying *SCN1A* DS *in vitro*.

### Transcriptional profiling reveals increased Medial Ganglionic Eminence specification and dis-regulation of developmental processes in SCN1A T1722N SSs

We profiled T1722N patient and control SSs by bulk RNA-sequencing and RT-qPCR at three stages of differentiation, DIV 35-40 (TP1, timepoint1), 75-80 (TP2, timepoint2) and 115-120 (TP3, timepoint3) (Figure 2).

**Figure 2.**
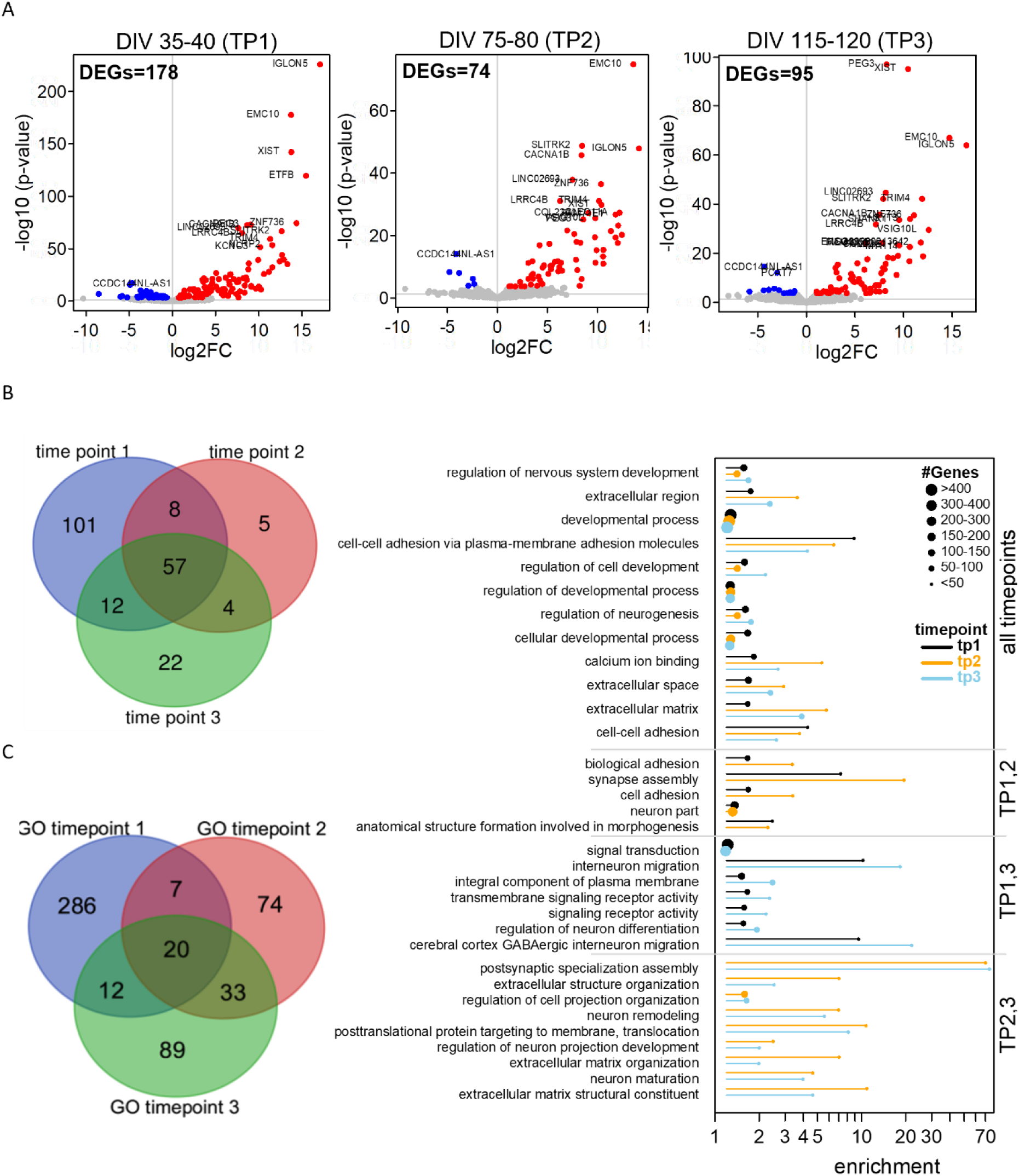
Bulk RNA-sequencing of *SCN1A* T1722N patient iPSC-derived SSs revealed major dis-regulation of developmental processes. (A) Volcano plots of significantly differentially expressed (FDR *q* < 0.05) genes (DEGs) in patient-derived SSs versus corresponding isogenic control at three differentiation timepoints (TP1, 2, 3). Significantly up- and down-regulated genes are shown in red and blue, respectively. (B) A total of 57 overlapping DEGs was found across the three timepoints grouping (**C**) into 20 significantly enriched (FDR *q* < 0.05) gene ontology (GO) terms listed on the right. The enrichment values are based on the standard hypergeometric distribution approach (54) with the circle size represented by *n*, the number of DEGs that were highly ranked (by small FDR *q*-value) and represented in each GO term.

A total of 178 differentially expressed genes (DEGs) was found at the first timepoint compared to 74 and 95 DEGs at timepoints 2 and 3, respectively. This finding indicated that a major *SCN1A* loss of function impact happens already at the initial stage of corticogenesis, despite *SCN1A* function being described later in development (Figure 2A) (23,24). Interestingly, among the DEGs found at the first timepoint, three crucial factors of the medial ganglionic eminence (MGE) specification, NKX2-1, LHX-6, and LHX-8 (25–29), displayed a significant increased expression in T1722N SSs (Figure 1B). The mRNA expression of NKX2-1 was also validated by RT-qPCR (Figure S2B).

A total of 57 overlapping DEGs across the three timepoints was identified and found to group within 20 Gene Ontology (GO) terms. Of these, the majority corelates with the cellular processes, with terms linked to regulation of neurodevelopment showing the most significant enrichment (>300 genes) (Figure 2C).

### Neurons from SCN1A T1775N spheroids have reduced intrinsic and synaptic activity

Whole-cell patch clamping of neurons from 300 µm slices were performed in both control- and patient- (clone 13) derived SSs at 4-5 months *in vitro*. While 40% of control-derived neurons (6 out of 15) were able to fire multiple action potentials (APs) in response to depolarizing current injections, patient-derived neurons were silent or only fired one small-amplitude AP (Figure 3A-B). This difference was not related to passive membrane properties (e.g. membrane capacitance, resting membrane potential or input resistance), which were similar between control and patient groups (Figure 3SA). As a LoF is expected for the *SCN1A* T1722N variant, we examined voltage-sensitive inward and outward currents in voltage clamp (Figure S3B). Although no significant difference was observed in peak inward current density between control and patient-derived neurons, the patient group showed a trend towards lower values with more neurons displaying no inward currents (Figure S3B left). As the peak inward current likely represents the effort from all sodium channels, the loss of NaV1.1 channel function in patient neurons may have been masked to some extent by the presence of other sodium channel subunits (see Figure 1C). There is an apparent correlation between peak inward current density and the ability to fire APs (Figure S3B left, black circles), which could explain the overall lack of excitability in patient neurons. On the other hand, both peak and steady-state outward current densities (representing current flowing through various voltage-gated potassium channels) trend towards lower values in the patient group (Figure S3B middle & right) to match the slightly lower peak inward current density.

**Figure 3.**
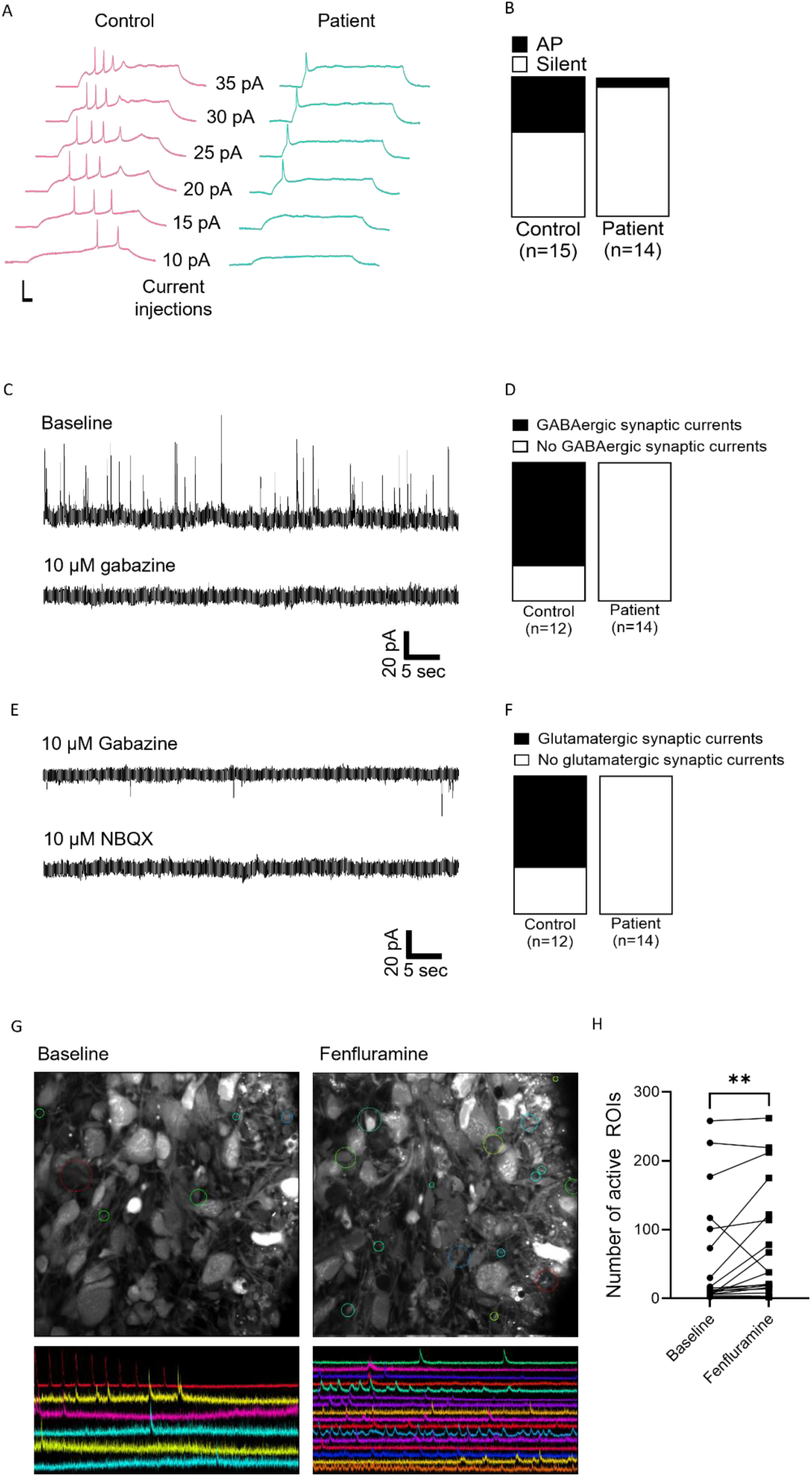
Whole-cell patch clamping of patient and isogenic control-derived SSs revealed reduced excitability and synaptic function in patient-derived neurons, which was rescued by acute treatment of 500 µM fenfluramine in two-photon calcium imaging. (**A**) Representative traces showing action potential (AP) firing in isogenic control (pink) and patient (teal) derived SSs. Scale bars, vertical = 50 mV and horizontal = 100 ms. (**B**) Proportion of cells capable of firing APs in control- (6) and patient- (1) derived SSs. (**C**) Spontaneous GABAergic synaptic currents recorded at a holding potential of +40 mV, before and after perfusion with 10 µM gabazine (GABAA receptor blocker). (**D**) Proportion of cells that received spontaneous GABAergic synaptic currents in control- (9) and patient- (0) derived SSs. (**E**) Spontaneous glutamatergic synaptic currents recorded at a holding potential of -70 mV, before and after perfusion with 10 µM NBQX (AMPA receptor blocker). (**F**) Proportion of cells that received spontaneous glutamatergic synaptic currents in control- (8) and patient- (0) derived SSs. (**G**) Representative images showing locations with Ca^2+^ transients before and after fenfluramine treatment in patient-derived SSs. Field of view = 275 µm x 275 µm. (H) Number of active ROIs before and after fenfluramine treatment. Mann-Whitney test, *p*<0.01.

We next investigated the effect of the T1722N variant on synaptic activity. The majority of control-derived neurons received spontaneous GABAergic (9 out of 12, Figure 3C-D) or glutamatergic (8 out of 12, Figure 3E-F) synaptic currents while 50% of recorded neurons received both types of synaptic currents (6 out of 12). Recording GABAergic synaptic currents from the ventral-forebrain derived spheroids was expected, while glutamatergic synaptic currents could be observed probably due to the presence of a small population of glutamatergic neurons (see VGLUT1 expression in Figure 1B). To distinguish the mode of neurotransmitter release, we examined the effect of Tetrodotoxin (TTX, 300 nM) on the spontaneous synaptic currents in a subset of cells. Most spontaneous GABAergic synaptic currents were blocked by TTX (Figure S3C), suggesting GABA release was evoked by spontaneous presynaptic AP firing. Interestingly, spontaneous glutamatergic synaptic currents were unaffected by TTX (Figure S3D), indicating that glutamate release was entirely due to spontaneous mechanisms at the presynaptic terminal. Additionally, we assessed whether NMDA receptor-mediated currents were present using the NMDAR antagonist D-AP5 and showed evidence of functional NMDAR in our SSs (Figure S3E-F). For example, a small proportion of spontaneous postsynaptic currents with the largest amplitudes disappearing (red oval in Figure S3E) and a shift of decay time from ∼50 ms to ∼5 ms (red arrows in Figure S3F) are indicative of NMDAR activity.

### Fenfluramine rescues the loss of neuronal function in patient-derived spheroids

As patch clamping studies revealed a clear functional phenotype of the patient-derived SSs, it would be valuable to see whether we can use this model to test ASM efficacy. To this end, we employed two-photon calcium imaging of intact patient-derived SSs and compared calcium transients observed before and after 500 µM fenfluramine perfusion. Encouragingly, fenfluramine increased the number of regions of interest (ROIs) displaying spontaneous calcium transients (Figure 3G-H), therefore potentially rescuing the loss of neuronal function in patient-derived SSs.

Many SSs also showed a characteristic spreading activity, with calcium transients originating from one cell or a confined location subsequently spreading to neighbouring cells on a time scale of tens of seconds (see examples in supplementary videos). While fenfluramine did not affect the number of recording sessions showing spreading activity in control-derived SSs (8 out of 16 at baseline, 8 out of 16 after fenfluramine treatment), it increased their number in patient-derived SSs by approximately 50% (from 7 to 10 out of 24). In a subset of control- and patient-derived SSs, we attempted to investigate the nature of different types of calcium transients (i.e. individual ROI activities not organized in any spatial pattern vs. spreading activity) using pharmacology. It is to our surprise that blocking GABAA receptors with gabazine had no apparent effects on any types of calcium transients. On the other hand, GABA appeared to enhance individual activity while preventing spreading activity. Interestingly, blocking AMPA receptors with NBQX enhanced both individual and spreading activity. These results suggest that spontaneous activity during early brain development has complex patterns and distinct mechanisms mediating each type of activity (data can be provided upon request).

## Discussion

As per many neurodevelopmental disorders, the pathophysiology of DS is still poorly understood and significant progress is hampered by the lack of robust models. However, while majority of other neurodevelopmental conditions presents with very heterogeneous genetics and clinical manifestations limiting the power of low-scaled *in vitro* studies, the etiology of DS primarily resides in pathogenic LoF variants of *SCN1A* and it correlates with homogeneous and defined clinical phenotypes. In this study we recruited a DS patient carrying the *SCN1A* T1722N mutation who manifests a severe and very defined clinical phenotype.

In rodent cerebral cortex, Nav1.1 is predominantly expressed at the axon initial segment of GABAergic interneurons and at a lower level in pyramidal neurons (22,30–34). Accordingly, DS epilepsy animal models have identified functional deficits in GABAergic interneurons, but not in pyramidal neurons (30,32,35). Despite some discrepancies in initial hiPSC-derived DS models (12,13), data shown so far represent compelling evidence of a functional decline in GABAergic neurons as the main driver of DS epileptogenesis (14,15,36).

Though the impact of *SCN1A* LoF variants on inhibitory neuron functionality has been widely investigated in human derived systems, little is known about the developmental pathomechanisms. This is likely due to the use of 2D cultures which were unable to recapitulate many aspects of *in vivo* human neurodevelopment, while the only 3D DS model lacked functional characterization (18). In addition, these studies compared patient-derived neurons with cells generated from either human embryonic stem cells or healthy control subjects. Therefore, in this study we exploited the unique property of recently developed 3D inhibitory differentiation protocol in recapitulating crucial aspects of cortical development and used this technology to derive subpullium-like spheroids from DS patient iPSCs and corrected control (19). The transcriptomic profile analysis of SSs revealed many DEGs, especially at the earliest timepoint. This was unexpected given the biological role of *SCN1A* described in later stages of neurodevelopmental and postnatal periods (23,24). As an important validation, our transcriptomic analysis showed that the expression balance of *SCN1A* and *SCN3A* during development is well recapitulated by our SS system with an opposite trend overtime (Figure 1D) (37). Therefore, although still preliminary, these data may suggest a non-canonical role of *SCN1A* in regulating human early corticogenesis. This was already reported for other *SCNxA* genes, e.g. *SCN3A* and *SCN9A,* which have been suggested to regulate neuronal development, such as neurite branching, neural migration and cerebral cortex folding, through AP-independent mechanisms, in addition to their established role in regulating sodium conductance during CNS development (38–40). In other forms of DEEs, recent iPSC-derived systems have revealed novel pathomechanisms in early stage of neurodevelopment driven by non-canonical expression and pathways of the candidate gene (41).

Interestingly, our transcriptomic analysis at the early timepoint reported a significant up-regulation of three crucial factors of the MGE specification, NKX2-1, LHX-6 and LHX-8, in patient SSs, which pointed to an increased ventralization and MGE specification in *SCN1A* LoF SSs. Of note, our finding was validated by RT-qPCR analysis and by comparing the expression of NKX2-1 across an additional control PSC line (data not shown) and mutant clone (clone 11) to further ensure the observed difference was not driven by an altered differentiation potential of the *SCN1A* control iPSC line or alternatively, by clone-specific abnormalities in the mutant cell line. Accordingly, by using a similar differentiation protocol, Zayat and colleagues showed a decreased expression of LHX-6 and other interneuron markers, DLX2 and SST, in DS patient-derived ventral forebrain-like organoids compared to no-isogenic control (18). This agrees with findings by others whereby mutations in risk candidate genes of DEEs (including Autism and LoF channelopathies) correlate with advanced interneuron development and excessive generation of GABAergic precursors and neurons (19,42–45).

In addition to revealing developmental disruptions, this 3D model shows the capability of recapitulating functional abnormalities of DS epilepsy. With fewer cells firing APs and receiving synaptic currents, DS patient-derived SSs displayed a clear LoF profile which was expected given the profound epileptic phenotype and defined clinical manifestation seen in the patient. These findings agree with functional profiling of 2D inhibitory neurons from DS patients carrying LoF *SCN1A* variants which showed reduced sodium conductance and AP firing as well as reduced GABAergic synaptic activity. However, the complete loss of synaptic activity including glutamatergic inputs in our patient-derived SSs is intriguing since pathogenic *SCN1A* variants are mostly expected to affect GABAergic neurons. In addition, glutamatergic synaptic transmission observed in the control-derived SSs was independent of presynaptic neuronal activity (see Figure S3D). One possible explanation for the global loss of synaptic activity could be related to GABA’s role in synaptogenesis during early brain development. It has been shown that GABA can induce *de novo* synaptogenesis of both GABAergic and glutamatergic synapses in a spatially precise manner (46). Therefore, the reduced intrinsic activity of GABAergic neurons in patient-derived SSs could be responsible for a global reduction in GABA release (see Figure S3C), leading to impaired synaptogenesis.

The functional deficit in the patient-derived SSs could be rescued to some extent by acute application of fenfluramine as assessed by two-photon calcium imaging (Figure 4G-H). Fenfluramine has been shown to offer significant improvement in DS patients, possibly via multiple mechanisms of action (47). For example, fenfluramine acts to increase serotonin release by inhibiting its reuptake while also inhibiting several classes of serotonin receptors such as 5HT-1D and 5HT-2A/C (48). These serotonin receptors are G-protein coupled receptors that have multiple downstream targets including potassium channels and Ca^2+^ signaling pathways (49), thus are capable of modulating neuronal excitability. Sigma 1 receptors, another main target of fenfluramine, can modulate various voltage- and ligand-gated ion channels at the postsynapse to indirectly influence synaptic function (47). It has also been demonstrated that serotonin can modulate GABAA receptors and inhibitory neurotransmission in hiPSC-derived neurons (50).

Several types of spontaneous Ca^2+^ transients were observed in our SSs, including individual ROIs and spreading activity, potentially recapitulating components of the network and oscillatory activities that occur during early brain development (51). In a subset of SSs, we attempted to dissect the nature of these distinct types of activity using pharmacology and showed that neither individual ROIs nor spreading activity were affected by blocking GABAA receptors. Intriguingly, GABA was seen to consistently increase individual neuronal activity which could be explained by GABA’s excitatory action during early development. It has been proposed that GABA acting on GABAA receptors first exerts excitatory effects which gradually switch to inhibitory effects owing to the changes in chloride gradient due to the developmental expression of several chloride transporters such as NKCC1 and KCC2 (52,53). While these chloride transporters are expressed at the mRNA level in our SSs, it is not possible to determine GABA’s excitatory vs. inhibitory effects based solely on this information. Protein expression and functional studies will be required to confirm our speculation. On the other hand, the spreading activities seem to be enhanced by blocking AMPA receptors. It has been shown that some cortical early network oscillations (cENOs) occurring at low frequency (∼0.01 Hz) are driven by NMDA receptors (51). It is possible that by blocking AMPA receptors, more glutamate was made available to NMDA receptors which upon activation caused the increase in network activity manifested as spreading activity in our SSs. Alternatively, the spreading activity could travel to neighboring cells via gap junctions as high expression of GJA1 (encodes connexin 43) was observed in our SSs at all three time points.

---

**Figure S1.**
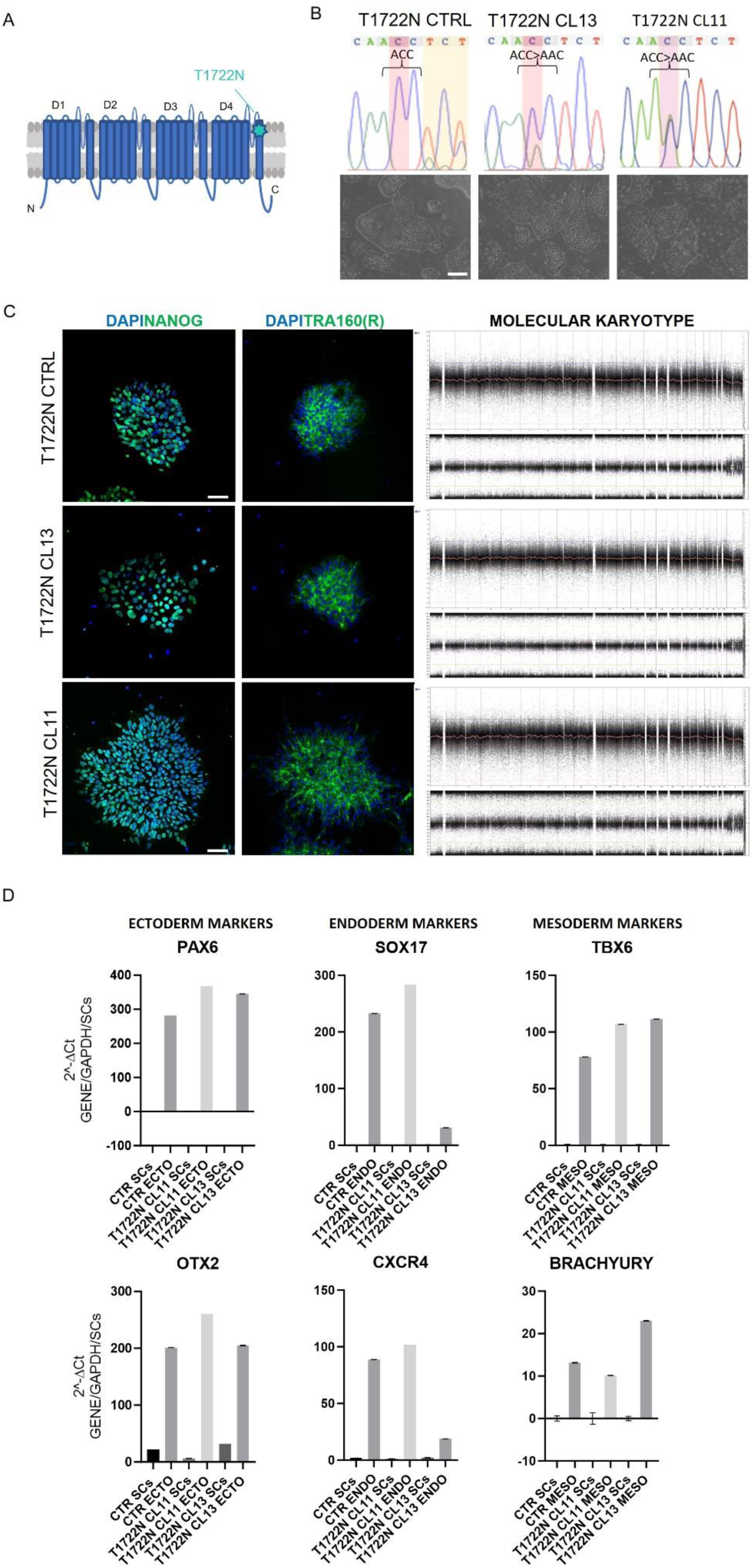
Characterization of *SCN1A* T1722N variant iPSCs. (**A**) Schematic of Nav1.1 protein showing the position of T1722N mutation (D, Domain; N, N-Terminal; C, C-Terminal). (**B**) Sanger sequencing of T1722N iPS cell lines and corresponding CRISPR-mediated corrected cell lines (top) showing normal morphology in representative bright field images (bottom). Location of the c.5165C>A mutation is highlighted in red. Heterozygous incorporation of the synonymous change (highlighted in yellow) is shown in the CTRL. (**C**) Immunoreactivity of patient iPSCs and corresponding isogenic control to pluripotency markers NANOG and TRA160R, and molecular karyotype assay. (**D**) Expression of ectoderm (PAX6 and OTX2), endoderm (SOX17 and CXCR4), and mesoderm (TBX6 and BRACHYURY) markers assessed by RT-qPCR. Data presented as mean ± SD of *n* = 3 technical replicates obtained from *n* = 1 biological replicate. Scale bar, 50 µm.

**Figure S2.**
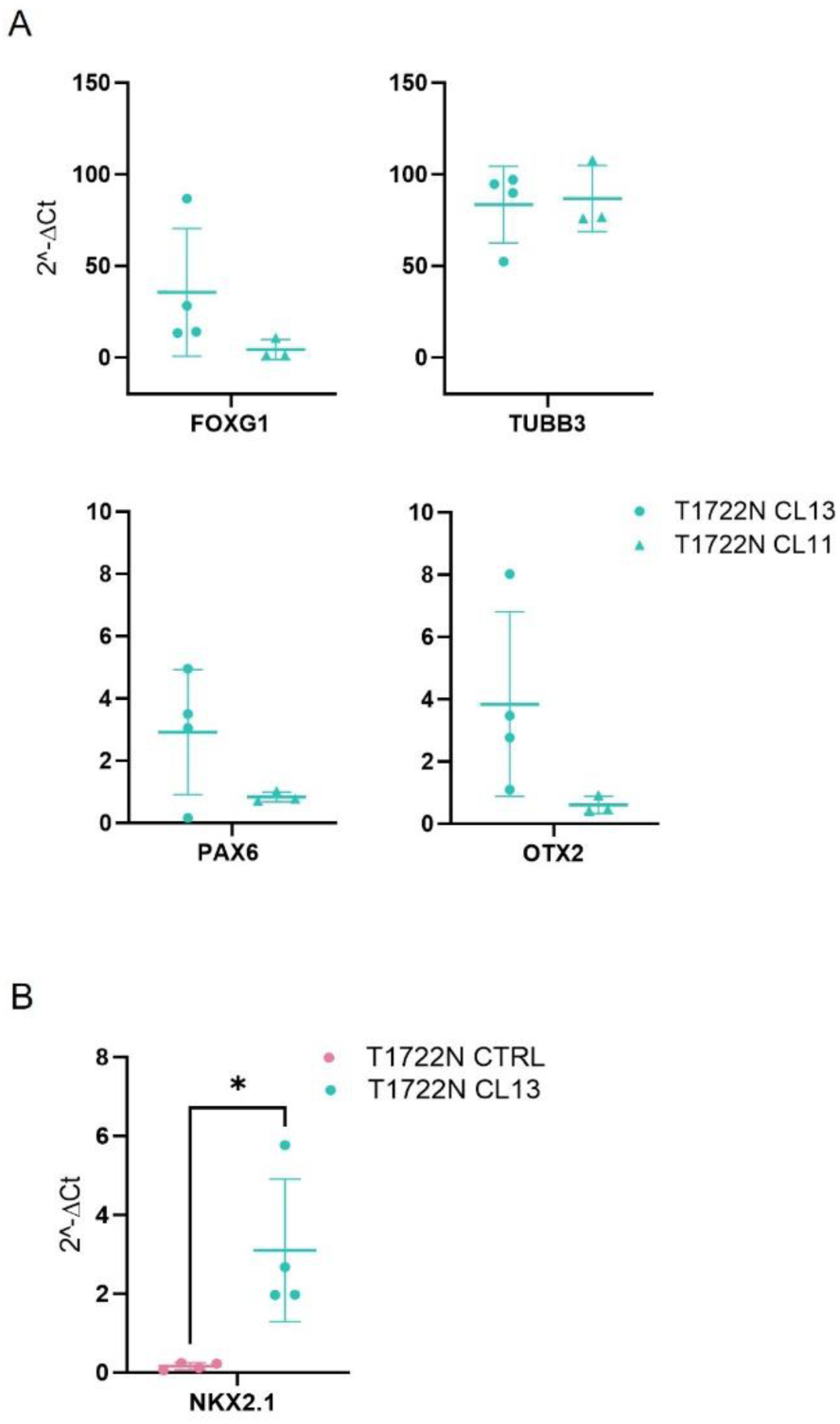
Characterization of *SCN1A* T1722N patient iPSC-derived spheroids. (**A**) Expression of neuronal (FOXG1 and TUBβIII) and dorsal-ventral axis (PAX6 and OTX2) markers in two distinct clones of patient iPSC-derived spheroids at DIV 35-40 assessed by RT-qPCR. (**B**) RT-qPCR of patient and control spheroids at DIV 35-40 showing the mRNA expression of ventral forebrain marker NKX2-1 (*p=0.0177). Data presented as mean ± SD obtained from *n* ≥ 3 biological replicates. The T-test was used to assess statistical significance (p<0.05).

**Figure S3.**
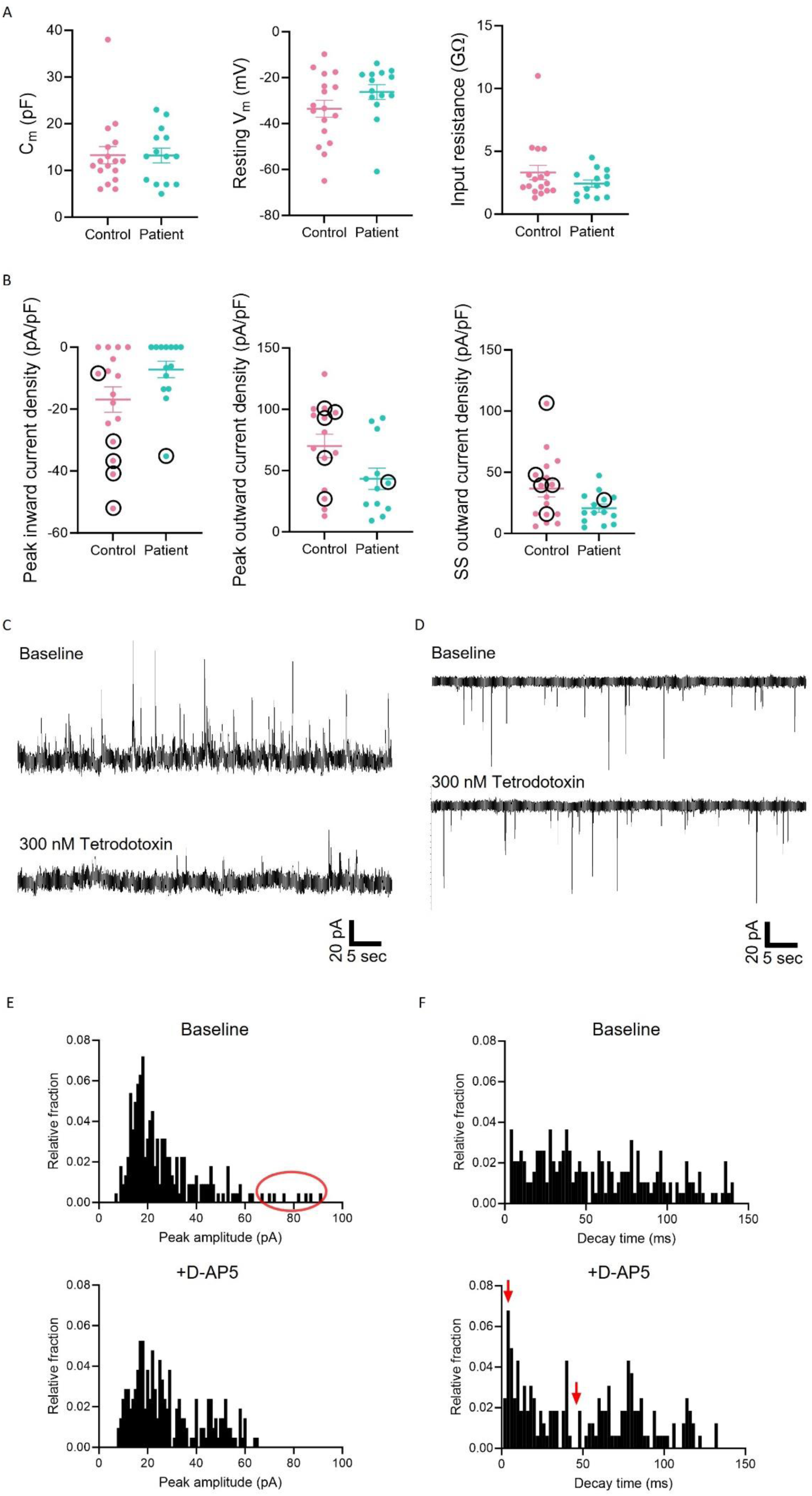
Additional electrophysiological characterisation of patient and control-derived SSs. (**A**) Passive membrane properties do not differ between neurons in patient or control-derived SSs. From left to right: membrane capacitance, resting membrane potential and input resistance. (**B**) Voltage-dependent inward and outward current densities are similar between neurons in patient or control-derived SSs. From left to right: peak inward current density measured for the largest inward current, peak outward current density measured at +40 mV, steady-state outward current density measured at +40 mV. Black circles indicate cells that were capable of firing action potentials with depolarising current injections from a holding potential of -70 mV. (**C**) Spontaneous GABAergic synaptic currents recorded at a holding potential of +40 mV, before and after perfusion with 300 nM tetrodotoxin (voltage-gated sodium channel blocker). (**D**) Spontaneous glutamatergic synaptic currents recorded at a holding potential of -70 mV, before and after perfusion with 300 nM tetrodotoxin. (**E**) Histogram showing relative fraction of spontaneous postsynaptic currents peak amplitude before and after perfusion of 25 µM D-AP5 (NMDAR antagonist) measured at +40 mV. Red oval indicates large synaptic events which disappear after D-AP5 perfusion. (**F**) Histogram showing relative fraction of spontaneous postsynaptic currents 90-10% decay time before and after perfusion of 25 µM D-AP5 (NMDAR antagonist) measured at +40 mV. Red arrows indicate disappearance of events with decay time around 50 ms and appearance of more events with decay time around 5 ms after D-AP5 perfusion. Mean±SEM. Mann-Whitney test.

## Notes

### Competing Interest Statement

The authors have declared no competing interest.

